# WU-CRISPR: characteristics of functional guide RNAs for the CRISPR/Cas9 system

**DOI:** 10.1101/026971

**Authors:** Nathan Wong, Weijun Liu, Xiaowei Wang

## Abstract

The CRISPR/Cas9 system has been rapidly adopted for genome editing. However, one major issue with this system is the lack of robust bioinformatics tools for design of single guide RNA (sgRNA), which determines the efficacy and specificity of genome editing. To address this pressing need, we analyze CRISPR RNA-seq data and identify many novel features that are characteristic of highly potent sgRNAs. These features are used to develop a bioinformatics tool for genome-wide design of sgRNAs with improved efficiency. These sgRNAs as well as the design tool are freely accessible via a web server, WU-CRISPR (http://crispr.wustl.edu).

## Background

The CRISPR/Cas9 system has been developed in recent years for genome editing, and it has been rapidly and widely adopted by the scientific community [1]. The RNA-guided enzyme Cas9 originates from the CRISPR-Cas adaptive bacterial immune system. CRISPRs (clustered regularly interspaced palindromic repeats) are short repeats interspaced with short sequences in bacteria genomes. CRISPR-encoded RNAs have been shown to serve as guides for the Cas protein complex to defend against viral infection or other types of horizontal gene transfer by cleaving foreign DNA [2-4]. Major progress has been made recently to modify the natural CRISPR/Cas9 process in bacteria for applications to mammalian genome editing [5, 6]. Compared with other genome editing methods, the CRISPR system is simpler and more efficient, and can be readily applied to a variety of experimental systems [7-11].

The natural CRISPR/Cas9 system in bacteria has two essential RNA components, mature CRISPR RNA (crRNA) and trans-activating crRNA (tracrRNA). These two RNAs have partial sequence complementarity and together form a well-defined two-RNA structure that directs Cas9 to target invading viral or plasmid DNA [2, 12]. Recent work indicates that it is feasible to engineer a single RNA chimera (single guide RNA, or sgRNA) by combining the sequences of both crRNA and tracrRNA [13]. The sgRNA is functionally equivalent to the crRNA/tracrRNA complex, but is much simpler as a research tool for mammalian genome editing. In a typical CRISPR study, an sgRNA is designed to have a guide sequence domain (designated as gRNA in our study) at the 5’-end, which is complementary to the target sequence. The rationally designed sgRNA is then used to guide the Cas9 protein to specific sites in the genome for targeted cleavage.

The gRNA domain of the sgRNA determines both the efficacy and specificity of the genome editing activities by Cas9. Given the critical roles of gRNA, multiple bioinformatics tools have been developed for rational design of gRNAs for the CRISPR/Cas9 system [14-17]. Experimental analysis indicates that Cas9-based genome editing could have wide spread off-target effects, resulting in a significant level of non-specific editing at other unintended genomic loci [14, 18-20]. Thus, most existing design tools have focused primarily on selection of gRNAs with improved specificity for genome targeting. However, more recent studies have demonstrated that the off-target effects of the CRISPR-Cas9 system is not as extensive as previously speculated, and random targeting of the noncoding regions in the genome has little functional consequences in general [21, 22]. Furthermore, novel experimental systems have been developed to improve the targeting specificity of CRISPR/Cas9 [23, 24]. Besides targeting specificity, another important aspect of bioinformatics design is to select gRNAs with high targeting potency. Individual gRNAs vary greatly in their efficacy to guide Cas9 for genome editing. Thus, the design of potent gRNAs is highly desired, as inefficient genome editing by Cas9 will inevitably lead to significant waste of resources at the experimental screening stage. The importance of gRNA efficacy has only been appreciated very recently, with multiple studies attempting to identify sequence features that are relevant to functionally active sgRNAs [21, 25-28]. For example, one recent study by Doench and colleagues analyzed 1841 randomly selected gRNAs and identified position-specific sequence features that are predictive of gRNA potency [21]. Similarly, CRISPRseek is a BioConductor package that also implements the Doench algorithm for potency prediction [29]. In our study, we reanalyzed this public dataset and identified many novel features that are characteristic of functional gRNAs. These selected features have been integrated into a bioinformatics algorithm for the design of gRNAs with high efficacy and specificity. A web server implementing this design algorithm has also been established.

## Results

In a recent study, Doench and colleagues analyzed 1841 sgRNAs to identify sequence features that are associated with CRISPR activities [21]. From that analysis, significant position-specific sequence features have been discovered. In particular, nucleotides adjacent to the protospacer adjacent motif (PAM), NGG in the target site are significantly depleted of C or T. In our study, this public dataset was systematically reanalyzed to identify other novel features that are predictive of CRISPR activity. To this end, we compared the most potent sgRNAs (top 20% in ranking) with the least potent sgRNAs (bottom 20%). By excluding sgRNAs with modest activities in this manner, distinct characteristics of functional sgRNAs can be more readily identified. The same strategy for feature selection has been proven to be effective in our previous study to characterize highly active siRNAs for target knockdown [30].

### Structural characteristics of functional sgRNAs

Previous studies have shown that structural accessibility plays an important role in RNA-guided target sequence recognition, such as by siRNA and microRNA [30-32]. Similarly, we hypothesized that structural characteristics of the sgRNA are important determinants of CRISPR activity. To this end, RNA secondary structures were calculated with RNAfold [33]. Overall secondary structure, self-folding free energy, and the accessibility of individual nucleotides in the structure were analyzed for each sgRNA. The sgRNA consists of two functional domains, the guide RNA (gRNA) sequence and trans-activating RNA (tracrRNA) sequence. The gRNA sequence consists of 20 nucleotides that pair perfectly to the targeted genomic sequence, thereby guiding the recruitment of the Cas9 protein to the target site; on the other hand, tracrRNA binds to Cas9 to form a functionally active RNA-protein (RNP) complex. As shown in Figure 1A, the tracrRNA region contains multiple well-defined structural motifs, which are important for interaction with Cas9 to form a functional RNP complex.

**Figure 1.**
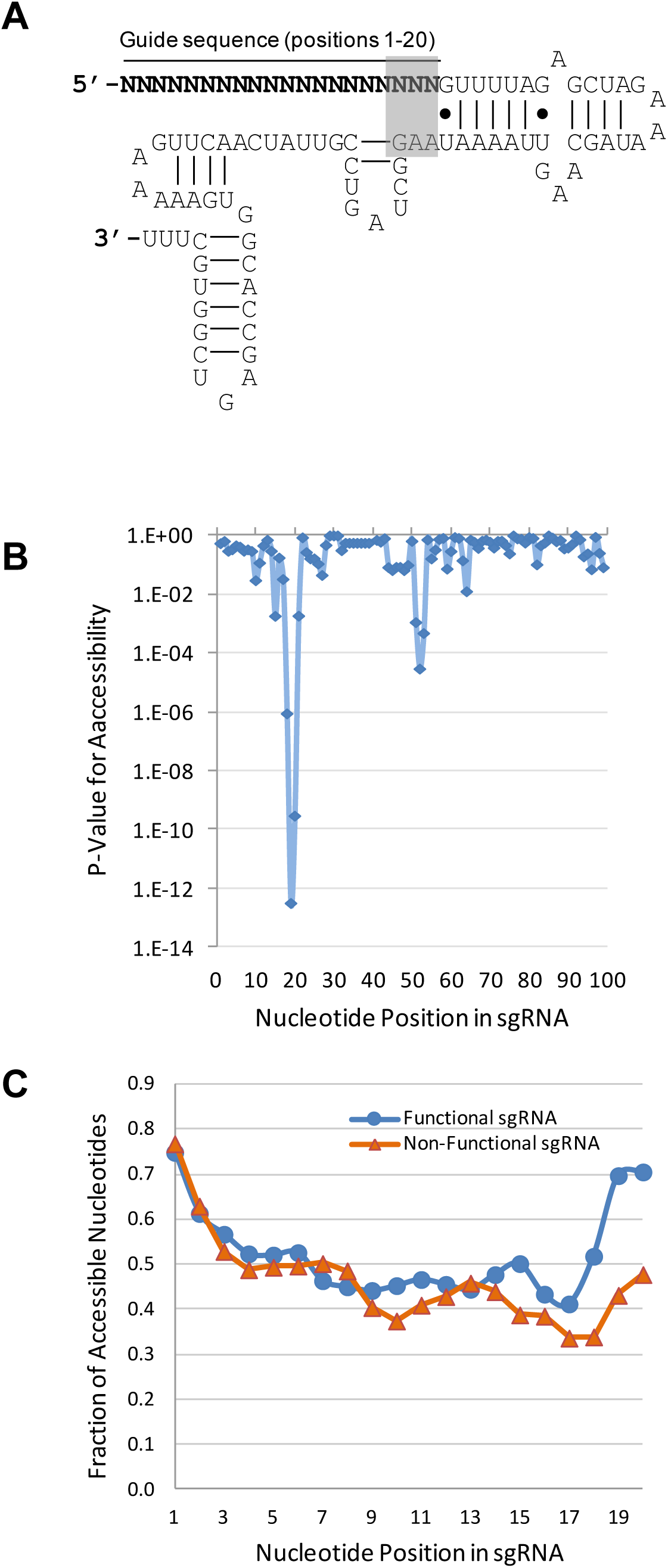
Structural characteristics of sgRNAs. **(A)** Secondary structure of the sgRNA. The 20-nucleotide guide sequence is complementary to the target sequence and resides at the 5’-end of the sgRNA. The highlighted nucleotides could potentially base pair, leading to an extended stem-loop structure. **(B)** Statistical significance of position-specific nucleotide accessibility of functional sgRNAs as compared with non-functional sgRNAs. **(C)** Comparison of position-specific nucleotide accessibilities between functional and non-functional sgRNAs.

Compared with non-functional sgRNAs, functional sgRNAs were significantly more accessible at certain nucleotide positions (Figure 1B&C). In particular, the most significant difference in accessibility involved nucleotides at positions 18-20, which constitute the 3’-end of the guide sequence (highlighted in Figure 1A). The 3’-end of the guide sequence, also known as the “seed region”, plays a critical role in recognition of target sequence. Thus, based on structural analysis, accessibility of the last three bases in the seed region was a prominent feature to differentiate functional sgRNAs from non-functional ones (Figure 1B). In addition, base accessibility in positions 51-53 was also significantly different. In the predicted structure of the sgRNA, nucleotides at positions 21-50 form a stable stem-loop secondary structure. From the survey of non-functional sgRNAs, nucleotides at positions 51-53 commonly paired with the end nucleotides of the guide sequence (positions 18-20), resulting in an extended stem-loop structure encompassing positions 18-53. Thus, decreased base accessibility at positions 51-53 was generally associated with decreased accessibility of the end of the seed region.

Furthermore, overall structural stability of the guide sequence alone (i.e. the gRNA domain residing positions 1-20) was evaluated with thermodynamics analysis. Specifically, the propensity of forming secondary structure was determined by calculating the self-folding free energy of the guide sequence. On average, non-functional guide sequences had significantly higher potential for self-folding than functional ones, with ΔG = -3.1 and -1.9, respectively (P = 6.7E-11, Figure 2A). Thus, the result from thermodynamic analysis also indicated that structural accessibility of the guide sequence was correlated with sgRNA functionality. In general, structural stability of the RNA can be approximated by the GC content of the sequence. Consistent with the free energy calculation, the guide sequence of non-functional sgRNAs had higher GC content on average as compared to functional sgRNAs (0.61 vs. 0.57, P = 2.1E-5). Furthermore, thermodynamic stability of the sgRNA / target sequence was evaluated. On average, non-functional guide sequences were predicted to form more stable RNA/DNA duplexes with the target sequence than functional ones, with ΔG = -17.2 and -15.7, respectively (P = 4.9E-10, Figure 2B). Thus, high duplex stability was a significant characteristic of non-functional sgRNAs.

**Figure 2.**
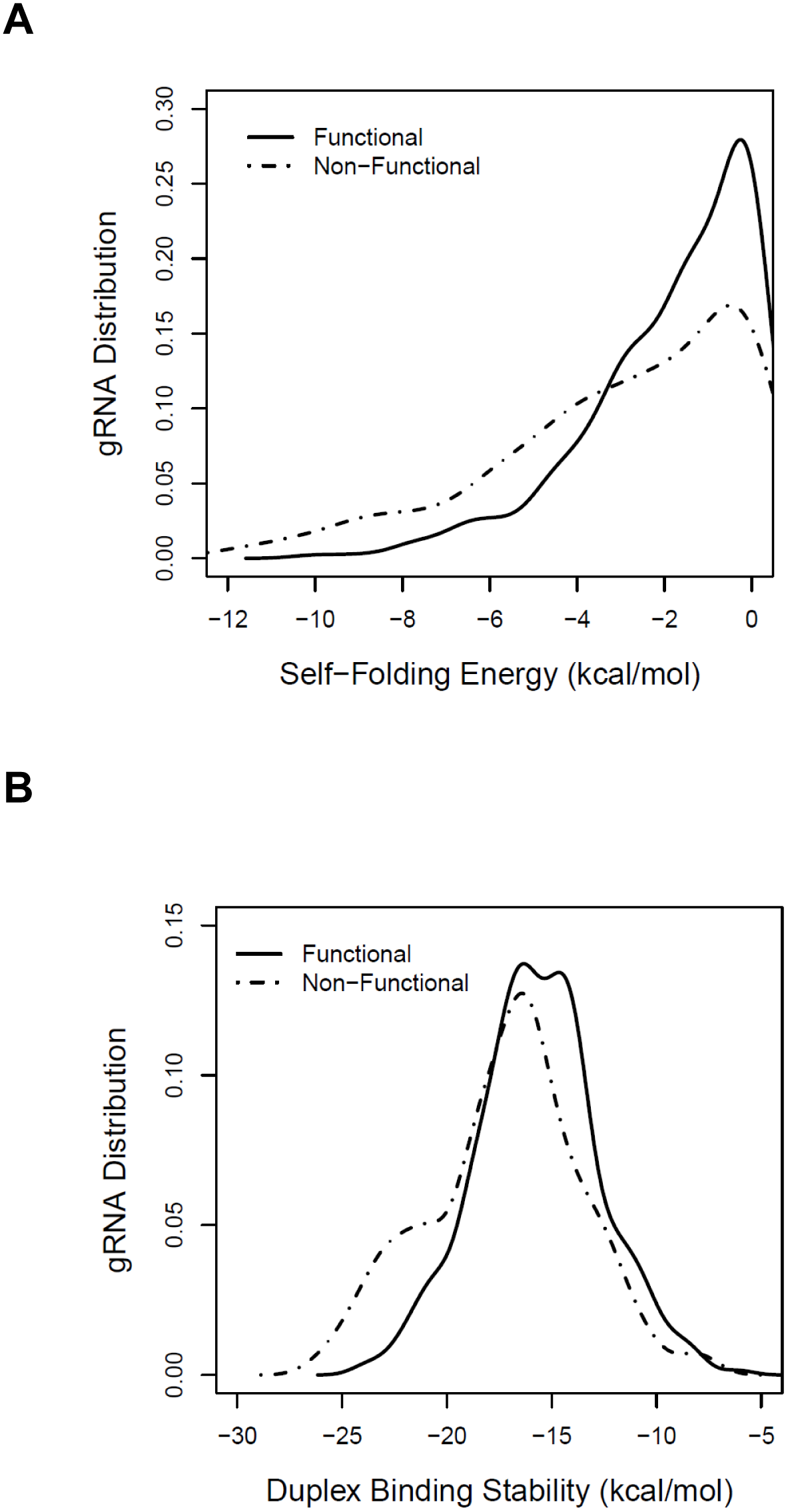
Thermodynamic properties of the guide sequence (gRNA). Functional and non-functional gRNAs were compared in the analysis. **(A)** Structural stability of the gRNA as evaluated by self-folding free energy (ΔG). **(B)** Structural stability of the gRNA/target sequence duplex as evaluated by free energy calculation.

### Sequence characteristics of functional sgRNAs

In addition to structural features describing the sgRNA, relevant sequence features of the guide sequence were also evaluated and are presented below.

#### Sequence motifs related to oligo synthesis or transcription

In most CRISPR applications, a 20-mer DNA oligo representing the guide sequence is cloned into an expression vector and expressed as the gRNA domain within the sgRNA. Thus, the efficiencies of both DNA oligo synthesis and subsequent transcription process are relevant to CRISPR activity. Repetitive bases (i.e. a stretch of contiguous same bases) could potentially be correlated with poor efficiency for DNA oligo synthesis. To assess this possibility, the distributions of repetitive bases in the guide sequence were compared between functional and non-functional gRNAs. Repetitive bases are defined as any of the following: five contiguous adenines, five contiguous cytosines, four contiguous guanines or four contiguous uracils. Overall, compared with non-functional gRNAs, functional gRNAs were significantly depleted of repetitive bases (5.4% vs. 22.8%, P = 1.3E-11). Among the four bases, four contiguous guanines (GGGG) were especially correlated with poor CRISPR activity. Previous work indicates that GGGG not only leads in poor yield for oligo synthesis, but also has the propensity to form a special secondary structure called guanine tetrad, which makes the guide sequence less accessible for target sequence recognition. Consistently, much fewer functional gRNAs were observed to contain the GGGG motif than non-functional ones (4.9% vs. 17.9%, P = 2.6E-8).

A stretch of contiguous uracils signals the end of transcription for RNA polymerase III that recognizes the U6 promoter. All gRNAs containing UUUU in the guide sequence had been preselected for exclusion from our analysis. Furthermore, recent work suggested that three repetitive uracils (UUU) in the seed region of the guide sequence could be responsible for decreased CRISPR activity [34]. Thus, a more stringent assessment was applied to evaluate the impact of potential transcription ending signal by searching for UUU in the last six bases of the gRNA. UUU was significantly absent in the seed region of functional gRNAs as compared to that in non-functional gRNAs (0.8% vs. 8.4%, P = 8.8E-7).

#### Overall nucleotide usage

Within the 20 n.t. gRNA sequence, the average counts for adenine were 4.6 and 3.3 for functional and non-functional gRNAs, respectively (P = 9.3E-18). In contrast, the usage of the other three bases (A, C or G) was only marginally correlated to CRISPR activity (Table 1, p-values in the range of 0.055 - 0.0019). The preference for adenine in functional gRNAs was not likely a mere reflection of overall preference for GC content as the uracil count was even lower in functional gRNAs than in non-functional ones (4.0 vs. 4.4). Overall usage of dinucleoside and trinucleoside were also examined and summarized in Table 1 and Supplementary Table S1, respectively. The most significant dinucleoside was GG (P = 2.3E-11) and the most significant trinucleoside was GGG (P = 4.9E-13). Both GG and GGG were significantly depleted in functional gRNAs, with enrichment ratios of 0.64 and 0.39, respectively.

**Table 1.** Significant base counts in functional gRNAs^*^

| Mono- or Di-<br>Nucleoside Count | Enrichment<br>Ratio | P-Value |
| --- | --- | --- |
| A | 1.39 | 9.3E-18 |
| U | 0.89 | 1.9E-03 |
| G | 0.92 | 6.2E-03 |
| C | 0.95 | 5.5E-02 |
| GG | 0.64 | 2.3E-11 |
| AG | 1.43 | 1.3E-09 |
| CA | 1.38 | 6.7E-09 |
| AC | 1.47 | 1.2E-08 |
| UU | 0.59 | 7.5E-08 |
| UA | 1.84 | 1.1E-07 |
| GC | 0.77 | 3.2E-06 |

\* The enrichment ratio was determined by comparing the average nucleoside counts of functional gRNAs to that of non-functional gRNAs. The p-values were calculated with Student's t-test.

#### Position-specific nucleotide composition

Base usage at individual positions was summarized and compared between functional and non-functional gRNAs (Supplementary Table S2). Consistent with previous findings [21], there was a strong bias against U and C at the end of functional gRNAs. Interestingly, a U or C at the end of the gRNA has a strong propensity to pair with AAG at positions 51-53 of the sgRNA, resulting in an extended stem-loop secondary structure (Figure 1A). Thus, the bias against U and C here was consistent with the structural analysis results, indicating the importance of free accessibility of the seed region for target recognition.

### Combining heterogeneous features for genome-wide prediction of sgRNA activity

Identified significant sgRNA features, including both structural and sequence features described above (summarized in Table S3), were combined and modeled in a support vector machine (SVM) framework. With these features, a computational algorithm was developed to predict the CRISPR activities. Similar to the sample selection strategy adopted in feature analysis, the most potent sgRNAs (top 20% in ranking) and the least potent sgRNAs (bottom 20%) were included in the SVM training process. The performance of the SVM model was validated by receiver operating characteristic (ROC) curve analysis. To reduce potential risk of overtraining, 10-fold cross-validation was performed in this ROC analysis. As shown in Figure 3A, the area under curve (AUC) was 0.92 for the SVM model. To further evaluate potential gene-specific bias in model performance, leave-one-gene-out cross-validation was performed. Specifically, experimental data from eight of the nine genes were used to train an SVM model while the data from the remaining gene were used for model testing in each iteration of the cross-validation process. The result of this gene-based cross-validation was similar to that of 10-fold cross-validation, with an AUC of 0.91. In contrast, a previous sgRNA prediction model based on the same training data had an average AUC of 0.76 from gene-based cross-validation [21]. Thus, our SVM prediction model could be used to differentiate functional sgRNAs from non-functional ones. In summary, cross-validation analysis indicated that our SVM model, which integrated both structural and sequence features, had robust performance at predicting sgRNA activities.

**Figure 3.**
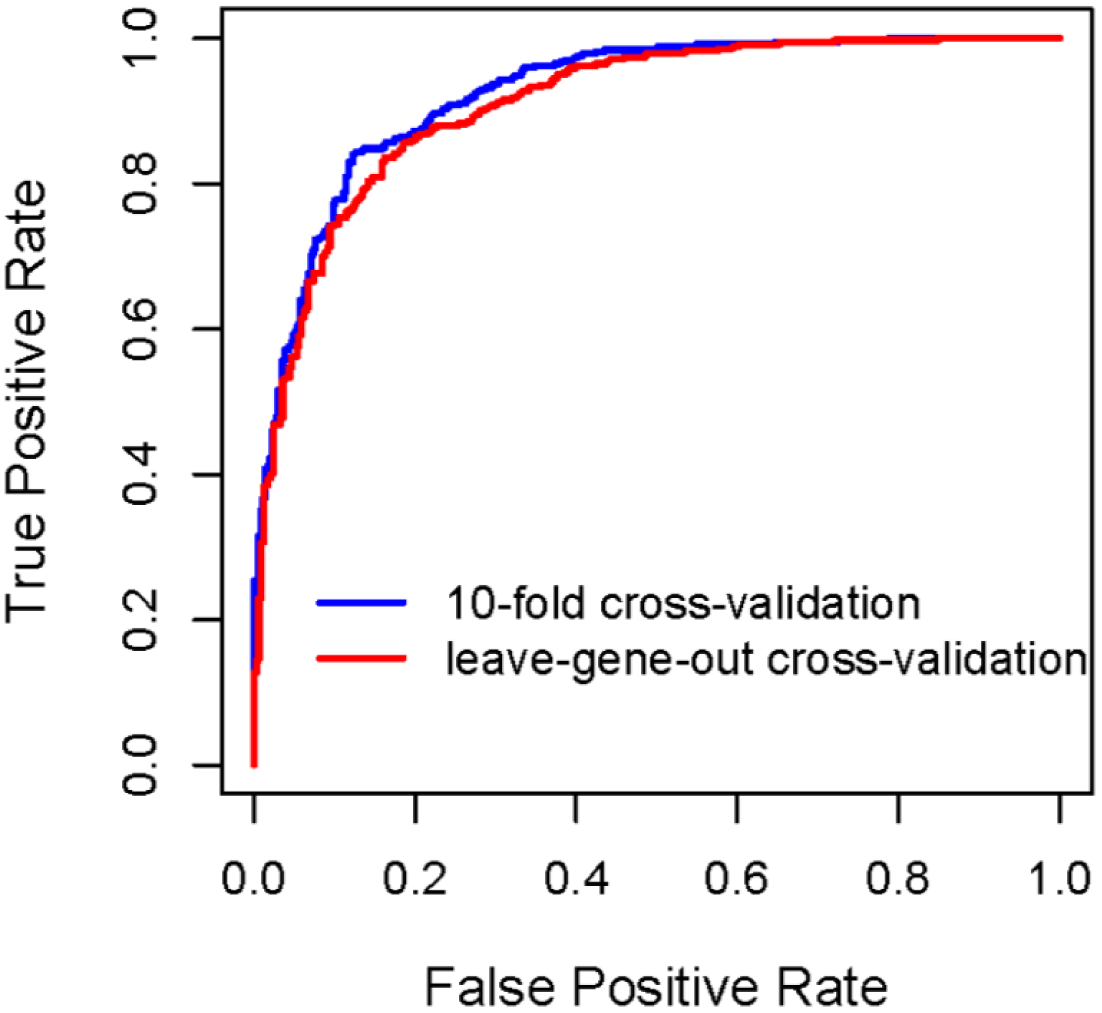
Evaluation of the gRNA prediction model by receiver operating characteristic (ROC) curves. Two cross-validation strategies were employed, 10-fold cross-validation and gene-based cross validation.

The SVM model was used to select functionally active sgRNAs for all known genes in the human and mouse genomes. To significantly speed up the selection process, a set of pre-filters were implemented to first quickly eliminate unpromising sgRNA candidates before evaluation by the SVM model. These pre-filters are summarized in Table 2. With these pre-filters, about 85% of non-functional sgRNAs were excluded while about 60% of functional sgRNAs were retained for further evaluation. Thus, application of the pre-filters led to a drastic reduction of non-functional sgRNAs while accompanied by only a moderate increase in false negative rate. By implementing these pre-filters before SVM modeling, a modified prediction model was constructed for genome-wide sgRNA design based on pre-screened training data.

**Table 2.** gRNA feature filters that were applied before the SVM modeling process.

| Filtered Features | Excluded Value | Enrichment Ratio for<br>Non-Functional gRNA |
| --- | --- | --- |
| gRNA folding ($\Delta G$) | < -8 kcal/mol | 15.8 |
| Duplex binding ($\Delta G$) | < -22 kcal/mol | 3.5 |
| GC content | > 80% | 30.7 |
| UUU in the seed region | True | 10.5 |
| Repetitive bases | True | 4.2 |
| Position 19 | U | 2.6 |
| Position 20 | C or U | 2.5 |

Free energy ( $\Delta G$ ) was calculated by RNAfold (23) for gRNA self-folding and by the Nearest Neighbor method (24) for binding stability of gRNA/target duplex.

The general applicability of the SVM model, which we named WU-CRISPR, was evaluated using an independent experimental dataset generated by Chari et al. [28]. In the Chari study, the knockout activities of 279 sgRNAs were determined experimentally by high-throughput sequencing and used to train a novel sgRNA design algorithm, sgRNAScorer. In our analysis, the activities of these sgRNAs were predicted with WU-CRISPR and correlated to experimental data. Furthermore, the performance of three other design tools, sgRNA Designer [21], SSC [27] and sgRNAScorer [28], were also evaluated using the Chari dataset. The Chari dataset was independent from WU-CRISPR, sgRNA Designer and SSC, but was used to train sgRNAScorer. Thus, ten-fold cross-validation results from sgRNAScorer (as presented in the Chari study) were included in our comparative analysis to reduce potential training bias. For each algorithm, top ranking sgRNAs were selected and their knockout activities were checked against the experimental results. Precision-recall curve analysis was performed to evaluate the prediction accuracy. Precision-recall curves are commonly used to evaluate prediction precision (proportion of true positives among all predicted positives) in relation to the recall rate (proportion of true positives among all positive samples). As shown in Figure 4, all four algorithms performed significantly better than random selection (113 functional sgRNAs among 279 tested sgRNAs, or 40.5% precision background). Among these algorithms, WU-CRISPR had the best performance at selecting functional sgRNAs. Specifically, all ten sgRNAs with the highest prediction scores by WU-CRISPR were experimentally confirmed to have high knockout activities. Similarly, among all 50 sgRNAs with the highest prediction scores by WU-CRISPR, 88% were experimentally validated for their high knockout activities.

**Figure 4.**
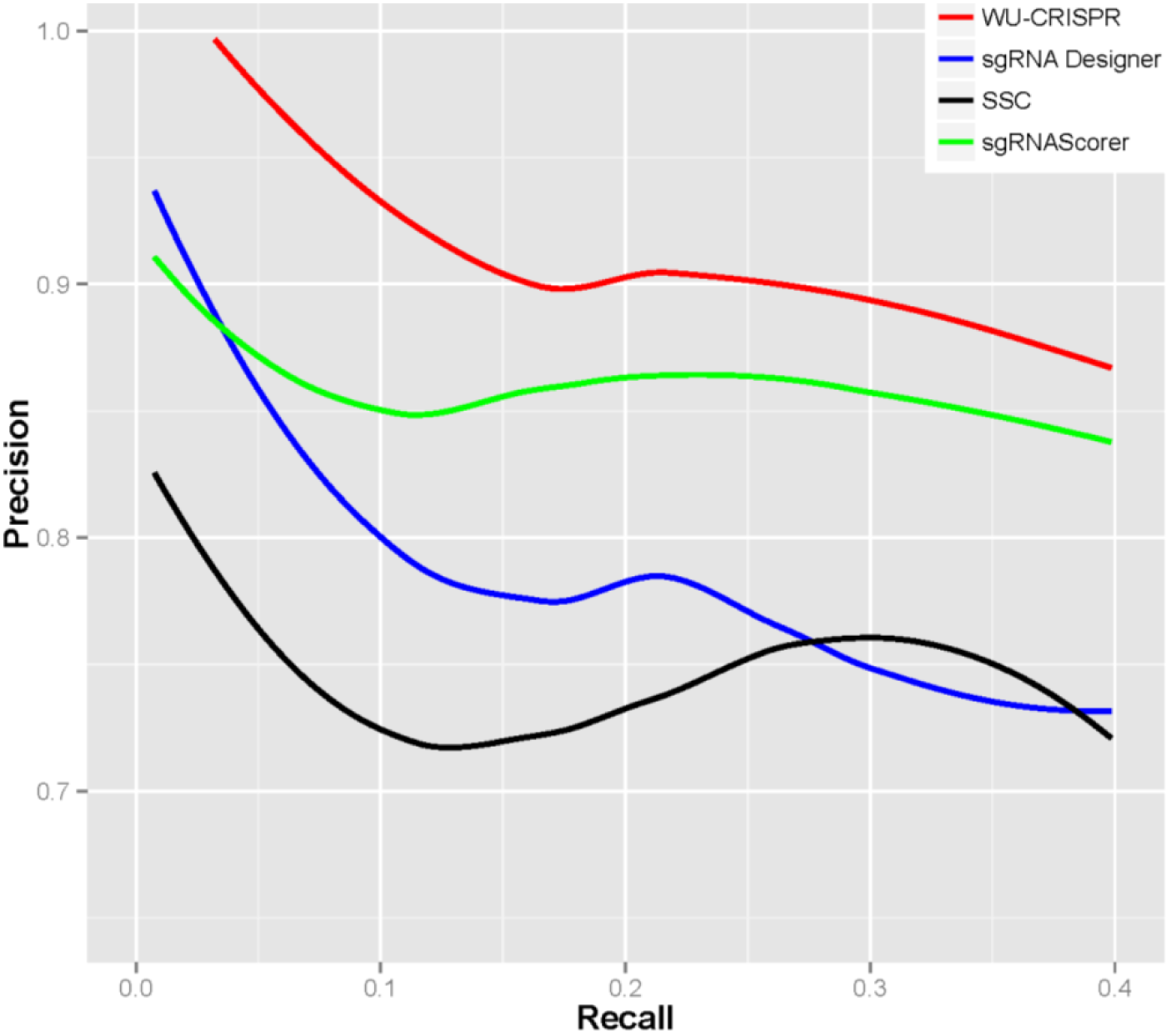
Validation of WU-CRISPR using independent experimental data. Precision-recall curves were constructed to evaluate the performance of WU-CRISPR and three other bioinformatics algorithms for sgRNA design.

Besides knockout efficacy, targeting specificity was also considered as an optional feature in the design pipeline. Targeting specificity of sgRNAs has been considered in previously published algorithms. However, existing algorithms search for potential off-target sites in the entire genome space. As the genome contains billions of nucleotides, sgRNA off-targeting is an unavoidable problem when all genomic regions are considered. Recent studies indicate that small-scale genomic alterations (insertions or deletions of less than 20 n.t.) induced by sgRNA had little functional consequence if the sites are within noncoding regions [21, 22]. Therefore, we decided to focus our off-targeting analysis exclusively on exon regions, including sequences from both protein-coding genes and other types of genes such as miRNAs and long noncoding RNAs. In this way, more stringent off-target filters could be implemented since a much smaller sequence space (as compared to the entire genome space) was searched.

Each gRNA candidate was compared to all known exon sequences in the genome. Recent experimental studies revealed that the 3’-end seed region of the gRNA is more relevant to off-targeting than the nucleotides residing in the 5’-end. Thus, a more stringent filter is applied to this PAM-proximal seed region. In our algorithm, a gRNA candidate was excluded if its seed sequence (3’-end 13 nucleotides) was found in any other unintended exon sequence preceding the PAM domain (NGG or NAG). Furthermore, BLAST sequence alignment was performed to identify and exclude 20 n.t. gRNA candidates that have over 85% similarity to any unintended sequence in the design space.

Using the established bioinformatics design pipeline to screen for both CRISPR efficacy and specificity, gRNA sequences were designed to target most known protein-coding genes in the genomes, including 18635 human and 20354 mouse genes, respectively. These gRNA sequences are freely accessible via a web server, WU-CRISPR (http://crispr.wustl.edu). In addition, a custom design interface was established for gRNA selection based on user-provided sequences.

## Discussion

In a short period of time, the CRISPR/Cas9 system has quickly become a major tool for editing of mammalian genomes. However, the rules governing the efficacy of CRISPR have not been well characterized and most users still design CRISPR assays by trial and error. This problem resembles a similar efficacy issue for RNAi studies ten years ago when the characteristics of functional siRNAs had not yet been well defined. As a result of significant advances in identifying the features that are characteristic of functional siRNAs, highly active siRNAs can be readily designed with bioinformatics tools, leading to a drastic saving in experimental resources. In the current study, we focused on identifying significant features that can be used to predict highly active sgRNAs. Specifically, we reanalyzed a public CRISPR dataset and discovered many novel features that are characteristic of functional sgRNAs. Previously, we and others have shown that both sequence and structural features of the siRNAs are important for RNAi knockdown activities [30]. Similarly, the knockout activities of CRISPR/Cas9 are also correlated to both sequence and structural features of the sgRNAs. By incorporating heterogeneous features in a prediction model, we have developed an improved bioinformatics design tool and implemented a web server, WU-CRISPR for genome-wide selection of gRNAs for the CRISPR/Cas9 system. The availability of this program may help to improve the efficiency of CRISPR assay design, leading to significant savings in experimental resources at subsequent screening stage.

## Methods

### Retrieval of public data for algorithm training

All gene sequences, including both exon and intron sequences, were downloaded from the UCSC Genome Browser [35]. Index files mapping transcript accessions to NCBI Gene IDs were downloaded from NCBI ftp site [36]. The Doench dataset for functional sgRNA screening was downloaded from the journal’s website [21]. In this published study, functional activities of 1841 sgRNAs were determined by flow cytometry. The Doench dataset was reanalyzed to identify novel features that are correlated to sgRNA efficacy.

### Computational tools and data analysis

LIBSVM was used to build computational models for sgRNA design (http://www.csie.ntu.edu.tw/~cjlin/libsvm/). For SVM analysis, a Radial Basis Function (RBF) was used for kernel transformation. Optimization of the RBF kernel parameters was done with grid search and cross-validation according to the recommended protocol by LIBSVM. RNA secondary structures and folding energies were calculated with RNAfold [33]. The predicted structures were examined at single-base resolution to determine whether individual nucleotides were base-paired or unpaired in the RNA structures. Statistical computing was performed with the R package (http://www.r-project.org/). Statistical significance (p-value) for individual features was calculated by comparing functional and non-functional gRNAs with Student’s t-test or χ^2^ test.

### Validation of WU-CRISPR with independent experimental data

The Chari dataset [28] was employed to independently evaluate the performance of WU-CRISPR. In the Chari study, the knockout activities of 279 sgRNAs designed for Cas9 (from *Streptococcus pyogenes*) were determined experimentally by high-throughput sequencing and used to train an sgRNA design algorithm, sgRNAScorer. In our comparative analysis, the Chari dataset was used to compare the performance of WU-CRISPR with three other public algorithms, including sgRNA Designer [21], SSC [27] and sgRNAScorer [28]. Ten-fold cross-validation results from sgRNAScorer was previously presented in the Chari study and included in this comparative analysis. The sgRNA Designer program was downloaded at http://www.broadinstitute.org/rnai/public/analysis-tools/sgrna-design; The SSC program was downloaded at http://sourceforge.net/projects/spacerscoringcrispr/. These stand-alone tools were used to predict sgRNA activities, and the prediction results were then compared to experimental data. Precision-recall curve analysis was done for algorithm comparison in R using the ROCR package, and plotted using the ggplot and stat_smooth functions in the ggplot2 package.

### Data Availability

The web server and stand-alone software package for gRNA design using the new design algorithm are distributed under the GNU General Public License and are available at http://crispr.wustl.edu. All sequencing data from the Doench study [21] and Chari study [28] can be retrieved from the NCBI Sequence Read Archive (accessions SRP048540 and SRP045596, respectively).

## Supplementary data

Supplementary data are available at the journal’s website.

## Authors’ contributions

XW designed the study. NW, WL and XW carried out research. XW and NW wrote the manuscript. All authors read and approved the final manuscript.

## Acknowledgements

We thank Matt Narens for technical assistance. We thank Raj Chari for providing cross-validation data as presented in [28]. This work was supported by the National Institutes of Health [R01GM089784 to X.W.].

